# The bacterial swarming factor SwrD belongs to a diverse family of proteins that form hexameric rings

**DOI:** 10.64898/2026.05.26.727983

**Authors:** Dhruva Nair, Kai Zhang, Angel Khala, Chunhao Li, Brian R. Crane

**Author notes:** For correspondence: **Brian R. Crane**, Weill Institute for Cell and Molecular Biology, Department of Chemistry and Chemical Biology Cornell University, Ithaca, NY 14853, USA.

## Abstract

Prokaryotic swarming is a collective surface-associated behavior that supports rapid colonization, biofilm formation, and host interactions. Swarming requires enhanced flagellar motor (FM) torque generation under high-load conditions. In many bacteria, this adaptation is achieved through increased stator unit recruitment and coordinated transcriptional regulation of motility genes. Genetic studies have implicated several relatively poorly characterized proteins in swarming, including SwrD, a component of the *fla-che* operon whose loss impairs torque generation in *Bacillus subtilis*. Intriguingly, SwrD is conserved in spirochetes that experience high viscous loads but do not exhibit canonical swarming, suggesting a broader functional role in FM regulation. Here, we structurally and biophysically characterize SwrD from *B. subtilis* and the spirochete *Borrelia burgdorferi*. In both cases, we show that SwrD assembles into a hexameric ring-like structure featuring a highly charged, disordered peripheral tail. Whereas SwrD displays structural similarities to periplasmic lipoproteins and DNA-binding proteins, lipidation regions and DNA interaction motifs of these other proteins are not conserved in SwrD. Deletion of *swrD* in *B. subtilis* results in pronounced swarming defects and altered expression of *fla-che* operon transcripts. In silico interaction analyses further identify the stator protein MotA as a high-confidence SwrD interactor, but this interaction could not be validated biochemically. Conserved SwrD residues mediate subunit contacts or locate to the hexamer periphery where they may mediate functionally important interactions.

## Introduction

Prokaryotes exhibit complex community behaviors such as biofilm formation and swarming motility that contribute to antibiotic resistance and virulence. These behaviors are often responses to diverse types of environmental stimuli that include surface encounters (mechanosensing) and the presence of nutrients and secondary metabolites secreted by members of the community (*1*). Sensing of external cues triggers transcriptional programs that promote metabolic and morphological adaptations to support the chosen lifestyle.

Cell swarming manifests as collective motion across moist solid surfaces and is found for a range of gram-positive and gram-negative bacteria, including *Escherichia coli*, *Pseudomonas aeruginosa*, *Proteus miribalis* and *Bacillus subtilis* (*2–4*). After reaching a specific density, cells become elongated and increase the number of their flagellar motors (FMs)(*5*). Their collective, synchronized motion allows rapid migration and colonization of the surface. Post swarm-phase, the community typically forms a biofilm (*6*). In the case of *B. subtilis,* colonization of the agriculturally relevant tomato root, swarming offers a symbiotic benefit: where the plants gain secreted microbicide and drought tolerance while the bacteria gains root exudates such as malic acid and other organic acids (*7*, *8*).

Swarmer cells must generate more FM torque to mitigate the high-cell density, viscous environment and increased load (*9–11*). As a result, cells increase the number of force transducing stator units, composed of MotA and MotB, and secrete surfactants. Genetic screens have demonstrated that several genes coding for proteins with unknown function cause swarming defects (*12*). For example, loss of SwrD (*ylzl* in *B. subtilis*), located on the *fla-che* motility operon, disrupts swarming likely through a mechanism attributed to decreased torque generation from the stator complex (*13*).

However, spirochetes require higher-torque generation by their periplasmic flagella to fulfill their ecological niche in highly viscous environments. Interestingly, some spirochetes such as the Lyme disease bacterium *Borrelia burgdorferi* do not present a swarmer phenotype (*14*), yet encode SwrD homologs. In spirochetes, the flagellar hook proteins FlgE are covalently cross-linked by a highly unusual post-translational modification called lysinoalanine (Lal) presumably allowing the FM to withstand higher-torque (*15*). The *fliL* gene codes for a protein that increases stator retention in the FM to also transduce higher torque (*16*, *17*). Thus, the SwrD genetic context and distinctive phylogenetic distribution warrant a deeper structural and functional characterization to further understanding of its potential contribution to torque generation in high-load contexts.

Herein, we biophysically characterized SwrD, from both *B. subtilis* (*Bs*), a bona fide swarming bacterium, and the non-canonical-swarming spirochete *Borrelia burgdorferi (Bb*). In both cases, SwrD forms a hexameric ‘donut’ structure with a highly-charged disordered tail localized on the periphery. Although SwrD resembles proteins involved in DNA binding, specific elements involved in DNA binding are not conserved by SwrD. In addition, SwrD shares sequence and structural homology to a family of periplasmic lipoproteins of unknown function whose subunit folds are circularly permuted relative to SwrD. We show that the *B. subtilis* 3A38 ΔswrD k/o strain is indeed swarming deficient and has aberrant expression levels of mRNAs that indicate a compensation in the regulation of the *fla-che* operon. *In silico* screens of potential protein-protein interactors reaffirm with high confidence that SwrD interacts with the stator component MotA.

## Results and Discussion

### SwrD is sparsely distributed amongst bacteria

To judge the phylogenetic distribution of SwrD we utilized the Pfam database taxonomy distribution of known SwrD annotations (PF06289) (*18*). SwrD is primarily found in Bacillota (∼62%), Actinomycetota (∼20%) and Spirochaetota (∼8%). Notably, SwrD is not found in some validated swarming genera such as Escherichia or Proteus and thus is not a strict requirement to elicit the phenotype. Furthermore, these three phyla exhibit differing flagellar motor configurations to the canonically described Pseudomonadota; Bacillota and Actinmycetoa are gram-positive with thicker extracellular peptidoglycan layers, while Spirochaetota contain endoflagella that reside in the periplasmic space.

Sequence analysis indicates that the SwrD protein consists of a conserved core domain of 48 residues with a species-specific extended C-terminal tail (CTT). The CTT ranges from 1-64 residues and has propensity for disorder (Fig S1).

To gain a clearer insight into SwrD structural conservation, an AlphaFold prediction of *B. subtilis* SwrD was used as input into the ConSurf server (*19*). The protein lacks conservation in the CTT and adjacent helix with the most highly conserved residues in the folded core: Asn15, Ile19, Glu23, Pro26, Asp27 and Thr28.

### Expression, purification and solution state properties of SwrD

To explore the structural properties of SwrD, we targeted homologues from the genetically tractable, agriculturally significant microbe *B. subtilis* and the Lyme disease causative agent *B. burgdorferi*. The two SwrD homologues share ∼46% sequence identity and represent two major clades that contain SwrD (Fig S2), while maintaining drastically different motor architectures (*20*). Hence, the comparison of the structures may provide insight into whether SwrD structurally contributes to a swarm/pseudo-swarmer phenotype in an evolutionary conserved manner.

SwrD expressed well, and circular dichroism analysis indicated that Bb SwrD has a larger coiled-coil component and greater thermal stability than Bb SwrD, with melting temperatures of 72 °C and 48 °C, respectively (Fig. S3). Size-exclusion chromatograph multi-angle light scattering (SEC-MALS) was used to investigate the solution-state properties of Bb and Bs SwrD. SEC-MALS revealed monodisperse peaks for both Bb SwrD and Bs SwrD, with the corresponding molecular weights (MWs) of 62 ± 2 kDa and 59 ± 5 kDa, respectively (Fig. S4). These MWs correspond to a hexameric oligomeric state for both proteins.

### SwrD molecular architecture

Bb and Bs SwrD, both crystallized as hexamers in space groups P2_1_2_1_2_1_ and P1, and the crystals produced atomic structures to 1.7 Å and 2.5 Å resolution, thereby corroborating the solution state quaternary structures (Fig. 1B,C). In addition, Bs SwrD packs crystallographically as a dimer of hexamers in a shifted ‘face-to-face’ configuration.

**Fig 1.**
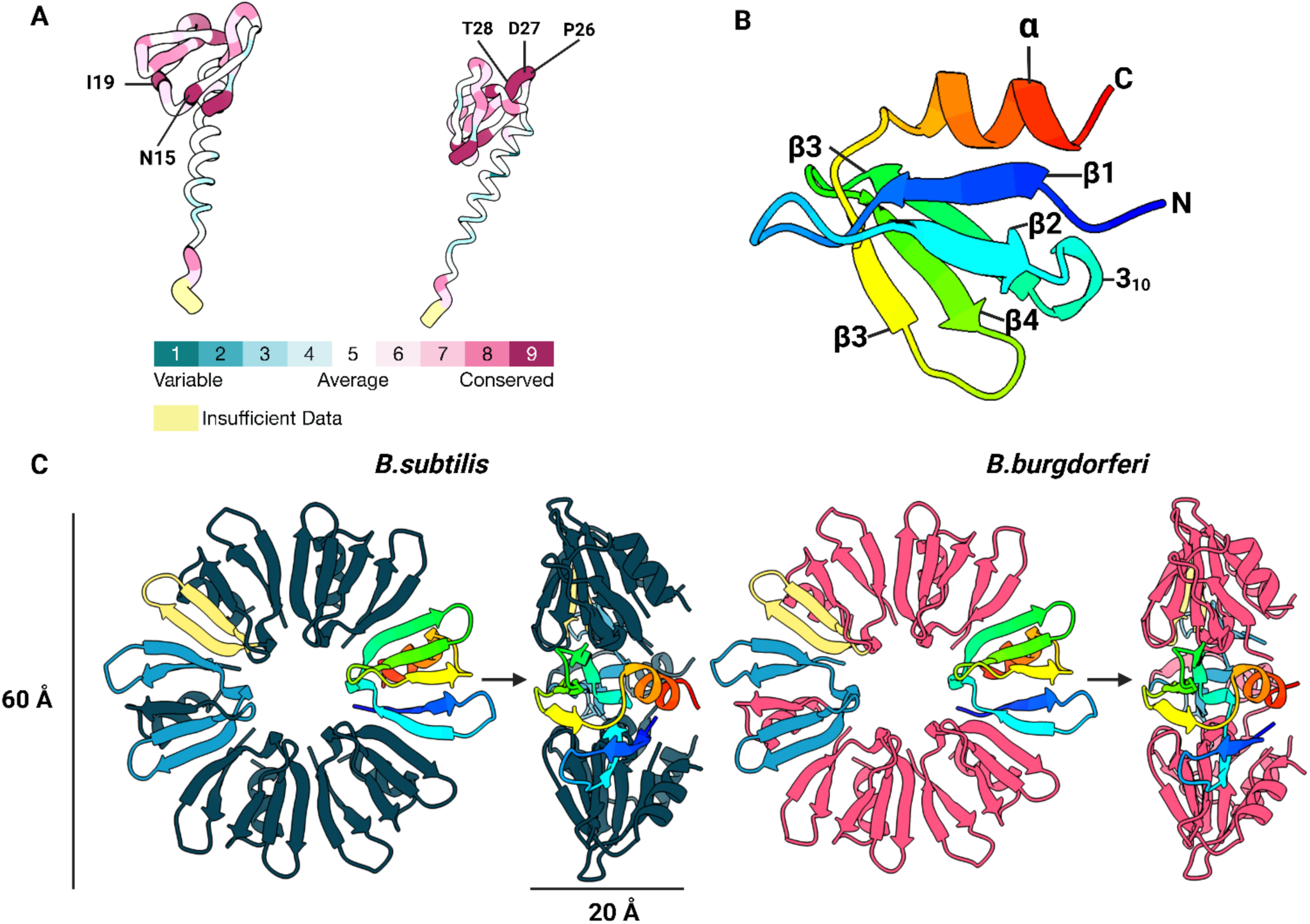
SwD conservation, secondary structure and molecular architecture. (A) Consurf performed on an AlphaFold model of monomeric SwrD, with highly conserved residues indicated. (B) Secondary structure of Bb SwrD colored blue to red (N to C). (C) Crystal structures of B.s SwrD and Bb SwrD, with a subunit colored as in B and the functional fold unit colored by chain in blue/yellow.

The subunits of each homologue share the same secondary structure and overall fold comprised of two anti-parallel β-strands that transition into a short 3_10_ helical loop followed by three anti-parallel β-strands and a terminal α-helix. Residues 52 to the c-termini for both SwrD homologues lack clear electron density, which is expected due to the disordered nature of the CTT (Fig. 1B,C,S1). The SwrD hexamers have an outer diameter of 60 Å with an inner diameter of 15 Å and a width of 25 Å. The β_1-2_ strands from one subunit form a continuous antiparallel 5-stranded β-sheet with β_3-5_ from the adjacent subunit (Fig. 1B,C). The C-terminal α-helices then pack against these interleaved cross-subunit β-sheets such that all of the helices lie on one face of the hexamer with the disordered C-termini directed toward the central pore. The central pore itself is composed of the six single-turn 3_10_ helices between β_1-2_ and β_3-5_ on one face of the donut and the six β_4_-β_5_ turns stacking them on the other side (Fig. 1B,C). Although the two SwrD homologues assume the same hexameric oligomeric state, the interfaces between subunits are mediated by different types of electrostatic and hydropathic interactions. In Bs SwrD electrostatic complementarity mates basic β_1-2_, and acidic B_3-4_ + α_1_; with the 3_10_ –helix and β_5_ bisecting the charged surfaces. Overall the structure maintains a neutral charge surface. On the other hand, Bb SwrD does not display the same disparate surface charges with β_1_ and α_1_ maintaining a neutral charge surface.

At the center of the donut, conserved Asn15 on the 3_10_ –helix hydrogen bonds with the peptide backbone of the 3_10_ helix of the adjacent subunit, as does His18 in Bb SwrD (Fig 2 A,D). Attempts to mutate Asn15 in Bs SwrD yielded insoluble protein, thereby highlighting its important contribution to SwrD stability. The oligomeric interface is further stabilized by salt-bridges between β_1_ -β_3_’ and β_4_/ β_5_ –β_3_. Bs SwrD maintains the additional Arg6-Asn21-Glu20 bridge to potentially compensate for the lack of His18 (Fig 2 C,D).

**Fig 2.**
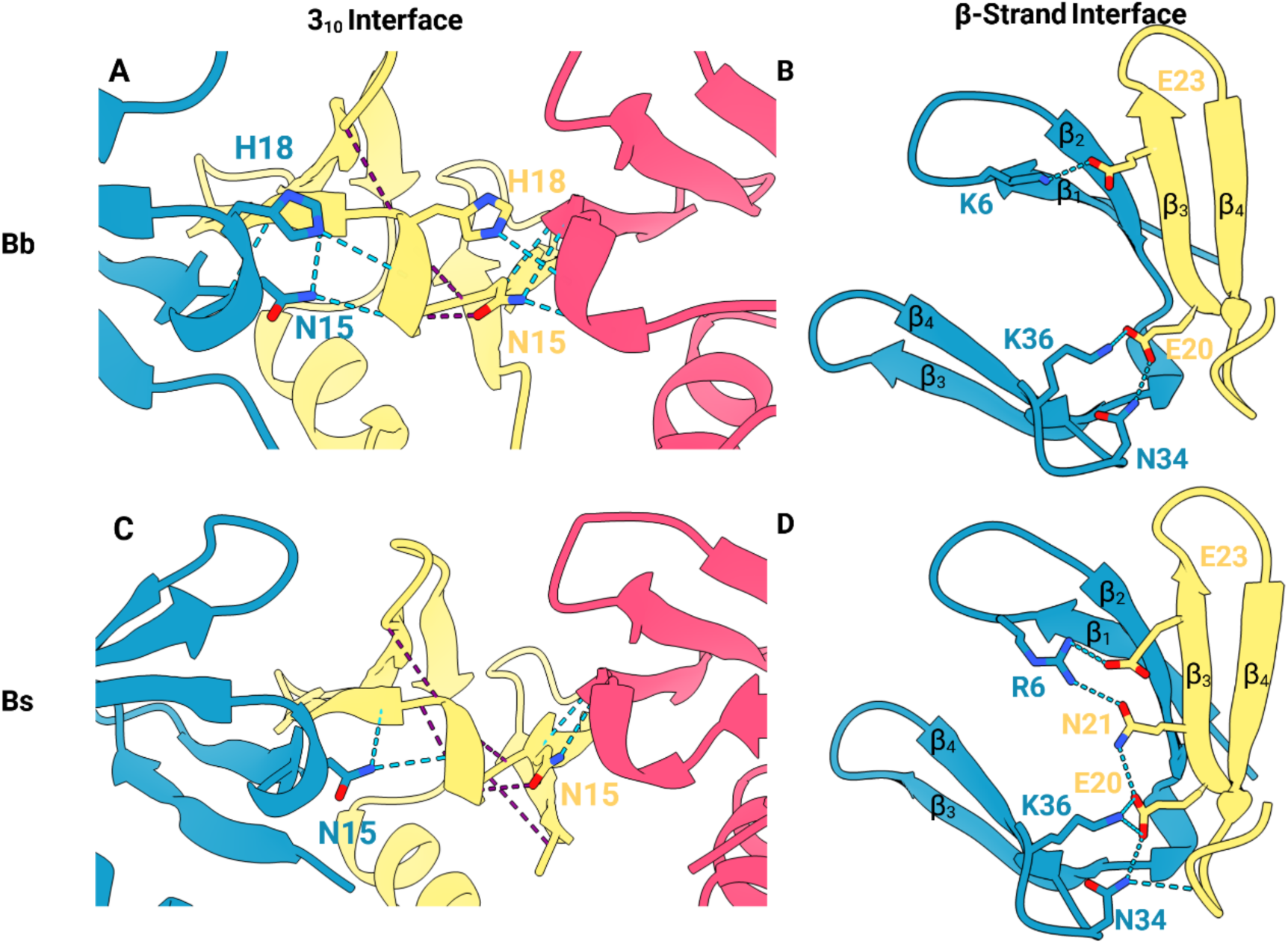
Key interfaces and interactions that stabilize the SwrD oligomer. Intermolecular hydrogen bonds that stabilize the hexamer shown in purple and intramolecular bonds shown in light blue. (A) Bb SwrD 3_10_ interface and (B) β-strand Interface. (C) Bb SwrD 3_10_ interface and (D) β-strand Interface

Structural conservation is particularly striking at three separate locations on SwrD i) β_3_, ii) Loop β_3_/ β_4_, and iii) the 3_10_ helix. β_3_ contains Glu20 and Glu23, that form salt bridges to the nearest neighbor subunit. Loop β_3_/ β_4_ contains the Pro Asp Thr motif that is surface accessible and lines the periphery of the donut and may be conserved for protein-protein interactions. Finally, the 3_10_ helix contains and Asn15-XXX-Leu/Ile19 motif, which is strongly conserved throughout all SwrDs, emphasizing the likely invariance of the hexameric oligomeric state of SwrD across organisms.

### Proteins structurally related to SwrD

Sequence similarity analysis predominantly identified SwrD homologs, whose genes also localized in flagellar operons. However, HHpred analysis also revealed several remote homologs from primary sequence similarity (*21*). The highest probability result was a member of the DUF903 family from the non-motile bacterium *Klebsiella pneumoniae* (PDB code: 5CAI). DUF903 proteins are predicted periplasmic lipoproteins of unknown function that include the *E. coli* protein YgdR (PDB:3FIF). The *K. pneumoniae* protein assumes a similar subunit fold and hexameric architecture compared to SwrD (Fig. 3), although *E. coli* YgdR itself crystallizes as a monomer. Identification of homology between SwrD and the DUF903 family is complicated by the fact that the DUF903 topology is circularly permutated relative to SwrD, with the N- and C-termini occurring at the SwrD β3/β4 loop, which contains the SwrD-conserved Pro-Asp-Thr motif (Fig. 3). The subunit interfaces of the respective hexamers also differ, generating somewhat perturbed overall structures that do not fully align (Fig. 3). The *K. pneumoniae* protein and close homologs conserve Asn64 (Asn15 in SwrD) that plays a critical role in stabilizing a hexamer, or some cases a heptamer (3BDU) in the central pore. In addition, the *K. pneumoniae* structure (PDB 5CAI) replaces Ile15 with Val68, which can be found in certain SwrDs. The protein families also differ in that YgdR homologs contain an N-terminal hydrophobic signal sequence followed by a lipidation signal (on a conserved Cys residue following the signal peptide), two features absent in SwrD, whereas SwrD contains the charged, disordered C-terminal tail, a feature absent from the DUF903 family.

**Fig 3.**
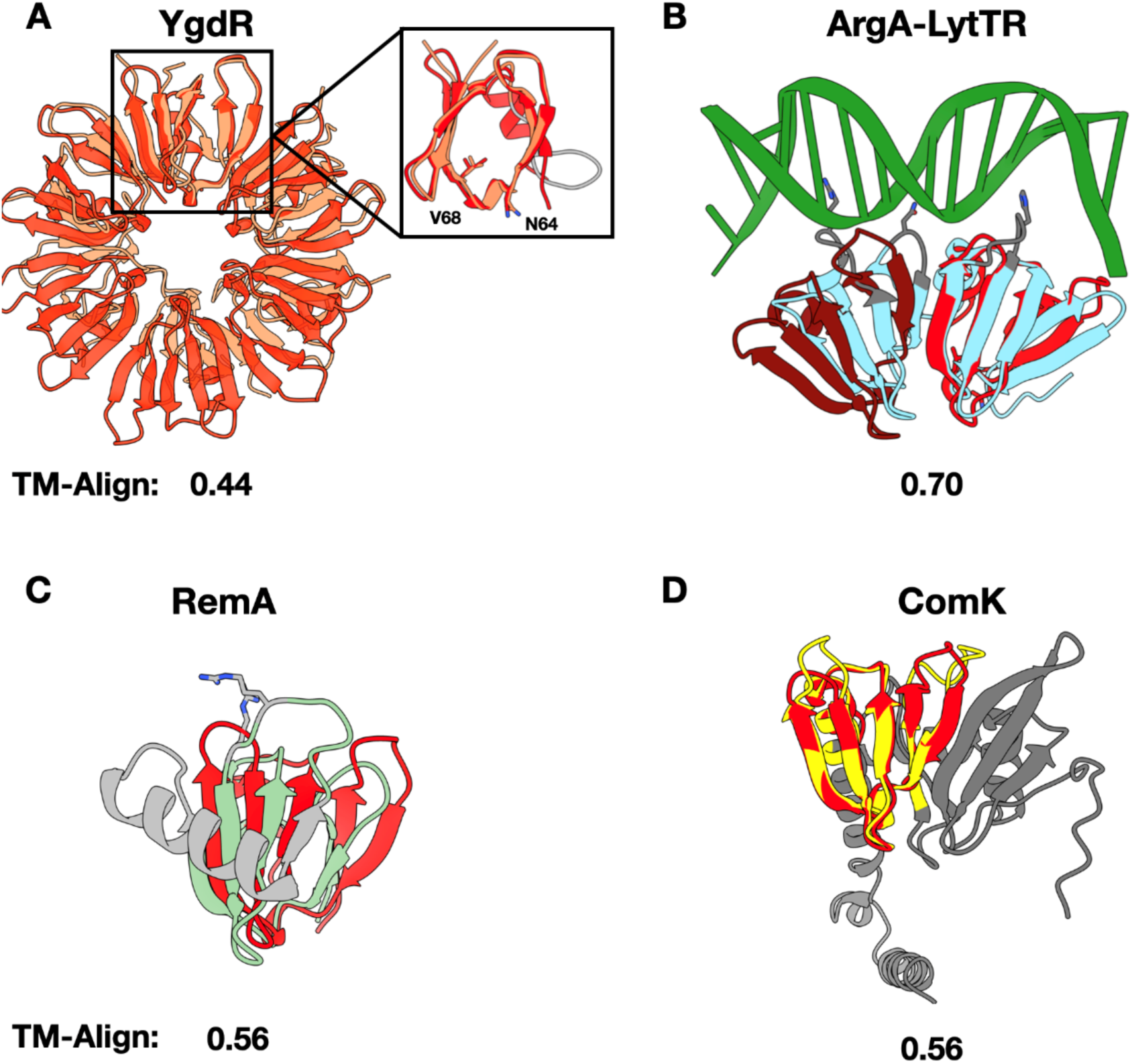
SwrD resembles diverse protein families. (A) Hexameric SwrD (orange-translucent) overlayed with hexameric DUF903/YgdR (salmon, PDB:5CAI). (B) SwrD dimer (red, marron) overlayed with ArgA-LytTR (light blue, PDB: 4XYO) bound to DNA (green, PDB: 4XYO), while the elements lacking in SwrD colored grey. (C) SwrD overlayed with RemA (sage, PDB: 7P1W) with elements lacking in SwrD colored grey. (D) SwrD overlayed with ComK (yellow, AF: A0A4R6B) with elements lacking in SwrD colored grey.

### SwrD is also reminiscent of small DNA/RNA interacting proteins

Three separate schemes (DALI, GT-align and FoldSeek) that prioritize 3D-structural similarity identified several protein families potentially related to SwrD: competence transcription factor (ComK, AFDB: A0A4R6BAQ9), the regulator of extracellular matrix A (RemA, PDB: 7P1W), and Accessory gene regulator A (AgrA-LytTR, PDB: 4XYQ). ComK,RemA and ArgA are DNA binding proteins involved in genetic competence, biofilm formation and quorum sensing, respectively (*22–24*). Although sequence similarities to SwrD are limited in all of these cases (11% - ComK, 20% - RemA, 29% - ArgA-LytTR; sequence identity relative to Bs SwrD), the subunit folds and oligomeric states among these proteins are remarkably similar, as illustrated by the 3D similarities among of β_1-5_ and 3_10_ (Fig. 3).

ComK acts as the master regulator of competency in *Bacillus*, which resembles a dimer of SwrD with a shortened helix (*25*). ComK is known to dimerize and tetramerize, although which quaternary structures are functional is currently unclear (*25*). RemA is a regulator of extracellular matrix formation that binds promiscuously to DNA sequences (*26*). RemA forms an octameric ring-like architecture like SwrD. The outer perimeter helical insertion between β3-β4 contains Arg51 and Arg53, which are responsible for DNA binding, which SwrD lacks. Bb SwrD aligns well with RemA (*Geobacillus thermodenitrificans* NG80-2), producing an RMSD of 0.76 Å in the regions excluding the β3-β4 helical insertion. The absence of the DNA binding motif in SwrD suggests that it does not function in the same manner as RemA. SwrD also shares fold similarity with the Helix-Turn-Helix (HTH) domain of LytTr from *Bacillus cereus* ATCC 10987 (Fig 3A). The HTH monomer of LytTR resembles a dimer of SwrD subunits (RMSD of 0.88 Å to one half of LytTR). It is unknown if LyTR forms a higher-order oligomer in solution. However, as SwrD lacks the DNA binding elements observed in LytTR, it is also unlikely that SwrD binds nucleic acids like LytTr. Nevertheless, the structural similarity between SwrD and these diverse proteins indicates that the β_1-2_-3_10_-β_3-5_ architecture acts as a robust modular scaffold that displays functionally specific structural elements at the periphery of the assembled multimers.

### A *swrD* defect alters swarming motility and shows associated gene-expression effects

To further explore the function of SwrD, we used the swarming-and competence-proficient *Bacillus subtilis* 3A38 (DK1042) strain to generate a SwrD knockout (*ΔswrD*) (*27*). Consistent with previous reports, deletion of *swrD* resulted in a pronounced swarming defect and a mucoid colony phenotype that has been shown to arise from the overproduction of poly-γ-glutamate (Fig. 4A) (*13*, *28*). Prior studies have shown that the *ΔswrD* swarming defect can be rescued by disruption of the anti-sigma factor FlgM, thereby increasing the activity of its cognate sigma factor, SigD (*13*). SigD increases the expression of late motility/chemotaxis/autolysin genes and of *flgM* (*29–31*). FlgM binds to SigD, and upon assembly of the hook-basal body, FlgM is secreted into the extracellular environment by the flagellar type III secretion system, relieving inhibition of SigD (*32*). Mutants in *flgM* that increase SigD activity and suppress *ΔswrD* are viewed to act through an offsetting mechanism, as *ΔswrD* deletion strains alone did not have altered SigD activity (*13*). In order to investigate similar mechanisms in the *ΔswrD* 3A38 strain we characterized transcript levels of *sigD* and its cognate anti-sigma factor *flgM*.

**Fig 4.**
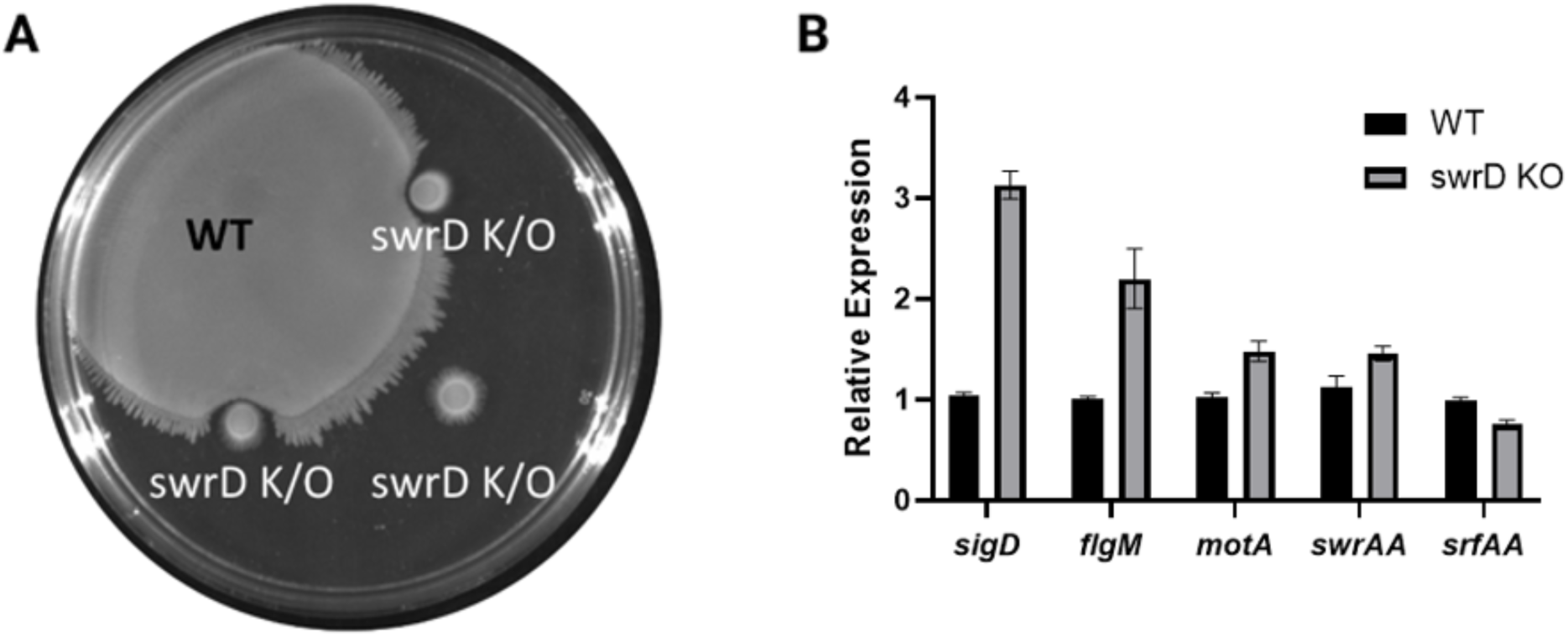
Swarming and mRNA expression of WT vs *swrD* k/o *B.s* 3A38. (A) Swarm expansion assay demonstrating that *swrD* k/o produces a swarming defect. (B) Relative expression levels of critical swarming genes between WT vs *swrD* k/o in planktonic high-cell density cultures. Statistical significance: *sigD*, *srfAA* and *motA*: P ≤ 0.0001, *flgM*: P ≤ 0.001 and *swrAA*: P ≤ 0.05.

We additionally examined expression of *motA* (a stator component whose overexpression increases motor torque in *ΔswrD*), *swrAA* (a regulator of swarming motility), and *srfAA*, which encodes a critical subunit of the surfactin synthetase complex (*13*, *33*, *34*). Because high-quality RNA could not be reliably obtained from swarm plates, high-density planktonic cultures were used as a proxy to approximate the cell densities encountered during swarm expansion.

Interestingly, *sigD* is expressed in ∼300% excess in *ΔswrD* vs WT. This change in *sigD* expression likely reflects compensation due to the mechanosensing potential of the flagella and its ability to effect the transcriptional profile of the cell (*35*, *36*). Disruption of the flagella is known to influence the global response regulator DegU, which can influence *sigD* expression (*37*, *38*). Unsurprisingly, the increased expression of *sigD* yielded a ∼200% increase of *flgM* in *ΔswrD* (Fig 4B). Increase in *flgM* expression could explain why SigD activity is not affected. SigD mediated increase in the transcription of *flgM* hence would prevent SigD from rescuing the *ΔswrD* defect (*13*). The mild ∼50% increase in the *sigD*-dependent *swrAA* and *motA* expression that are not enough to rescue the swarming defect can also be explained by relatively high *sigD* levels. Nonetheless, this dysregulation of *sigD* and its related genes is surprising in planktonic cells, and could imply that disruption of *swrD* expression may indirectly affect regulation of the *fla-che* operon. Regardless, the change in *sigD* levels do not explain the swarming defect of the *ΔswrD* mutant strain and likely reflect an indirect response to the motility defect (*13*).

### Structural implications of the central pore for oligomerization

As SwrD adopts a RemA-like fold and assembly state we sought to mimic mutagenesis studies of RemA that tested the determinants of the oligomeric state (*26*). Specifically, substitution of RemA Arg18 to Trp at the octamer central pore changed RemA from an octamer to a heptamer. The SwrD pore residue Tyr17 aligns well structurally with the critical oligomerization determining residue Arg18 (RemA, octamer) and Arg18Trp (RemA, heptamer). Hence, we generated residue substitutions to target the oligomerization of Bb SwrD: Tyr17Arg, Tyr17Trp and Pro16His:Tyr17His (Fig 5). Surprisingly, SEC-MALS and x-ray crystallography indicated that there was no change in oligomeric state caused by any of these residue substitutions (Fig. S4C,D). Both variants crystalized in P2_1_2_1_2_1_ (same as WT), and produced nearly identical structures to 1.8 Å (Tyr17Trp) and 1.9 Å (Tyr17Arg) resolution. To further explore the tolerance of the Bb SwrD 3_10_ –helix to residue substitution, the solvent accessible residues (Pro16 and Tyr17) were changed to His (Pro16His and Tyr17His) and a crystal structure of the variant was determined in the I4_1_22 space group. The variant was stable and formed a hexamer, similar to all of the other studied variants and WT, further indicating the ability of SwrD to act as a stable scaffold, one that could be potentially exploited for protein engineering applications.

**Fig 5.**
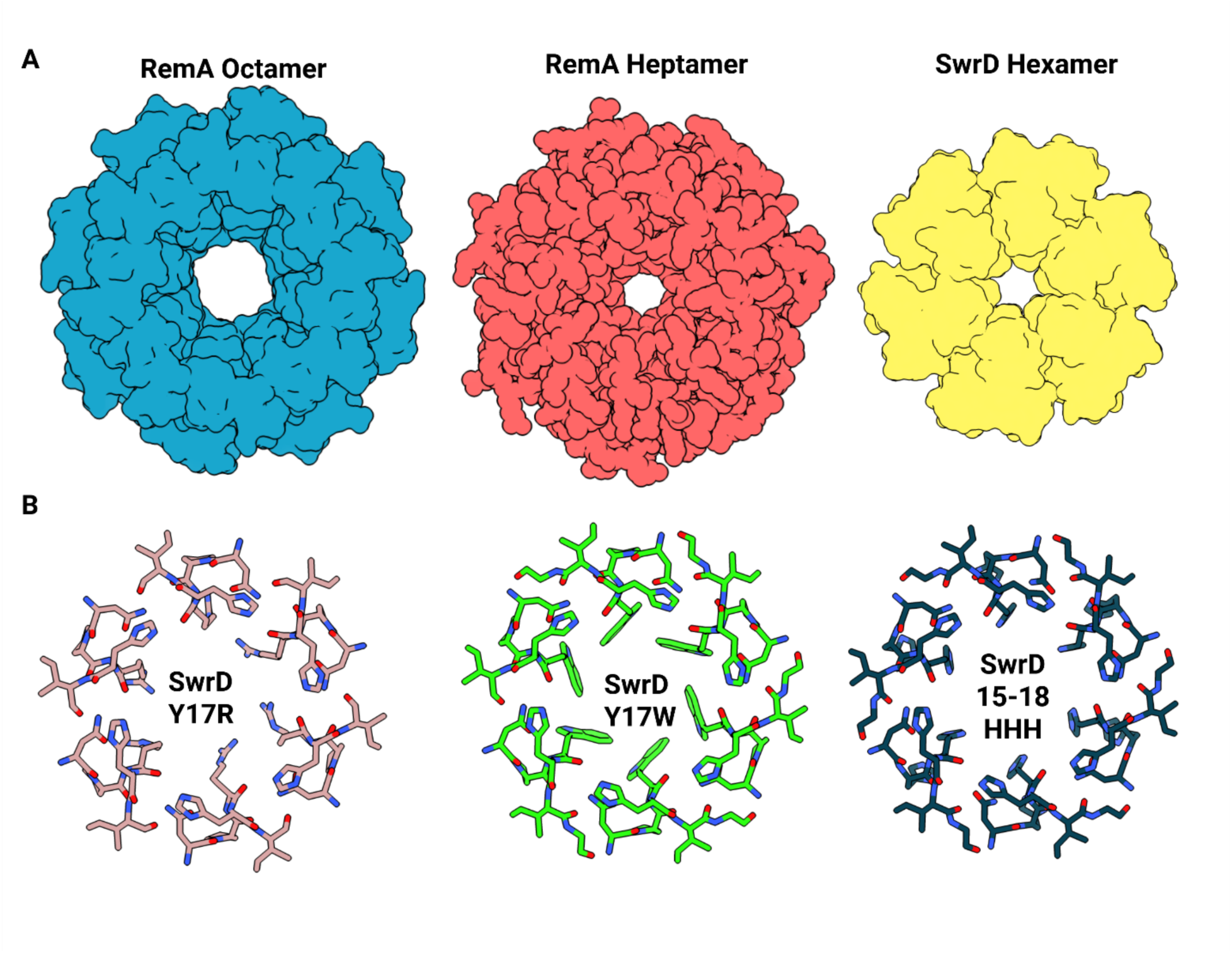
SwrD is a highly mutatable scaffold. (A) Space fill representation of RemA Octomer (PDB: 7BM2, blue), RemA Heptamer (PDB:7BME, red) and Bb SwrD (yellow). (B) Bb SwrD variants with 3_10_ helix residues shown.

### In silico SwrD interaction studies and attempted validation

To identify potential interactors of SwrD in *B. subtilis* we applied AlphaPulldown to predict multimers between SwrD and all genes associated with swarming (98 candidates) from the *Subti*Wiki database, with the SwrD – SwrD prediction as the internal benchmark (*39*). 20 candidates scored above 0.20 iPTM_PTM score threshold, with CheB, MotA, PgcA and CheD scoring above the SwrD to SwrD prediction (0.44), which given the known oligomeric state of SwrD, served as a positive control (Fig 6A,B).

**Fig 6.**
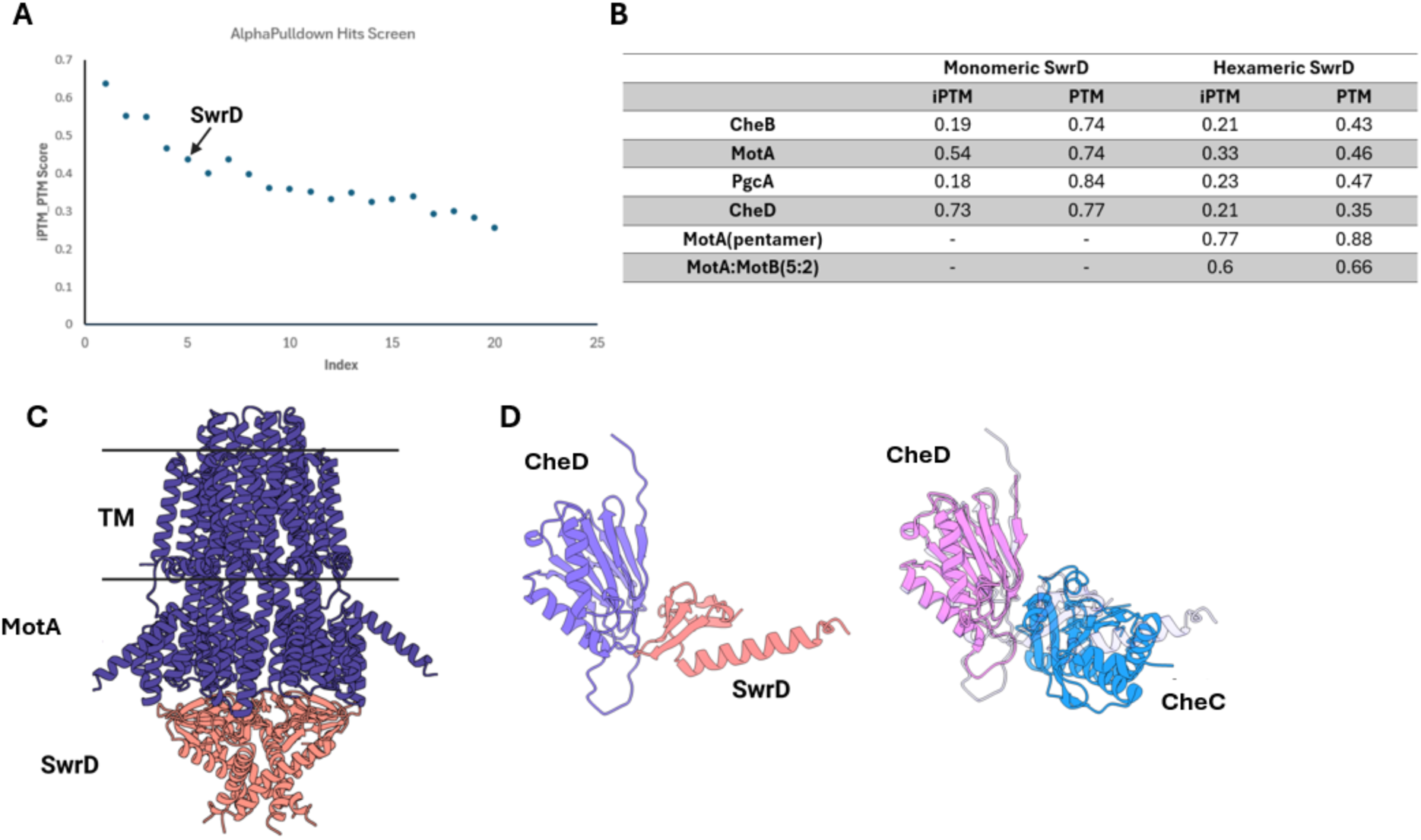
In silico pulldowns and multimer predictions show that SwrD potentially interacts with MotA and CheD. (A) iPTM_PTM scores of top hits plotted in descent, with SwrD highlighted. (B) Alphafold3 predictions of top hits in functional oligomeric states. (C) MotA (5 chains, purple):SwrD (6 chains, salmon) prediction. (D) CheD (light purple) and SwrD (salmon) interaction prediction (left), overlayed with CheD:CheC (PDB: 2F9Z, right).

AF3 predictions of these hits indicated that monomeric SwrD binds to the receptor deamidase CheD with an iPTM confidence of 0.73, on par with pentameric MotA, 0.77. Overlaying the SwrD-CheD prediction with the published crystal structure of the complex between CheD and the regulator CheC, suggested that SwrD may interact with CheD at the same interface as does CheC (Fig 6 B-D) (*40*). However, caution is warranted because the interaction predicts a monomeric form of SwrD, whereas SwrD does not form a monomer in solution under any conditions tested.

Intriguingly, the MotA-SwrD predictions locate SwrD to the MotA cytoplasmic region. Although this complex has a symmetry mismatch, the MotA central pore lines with the SwrD pore, with a subset of SwrDs conserved Arg6 and Asp27 lining the interface. We attempted to validate both the CheD and MotA interactions by in vitro interaction assays. In the first case, we produced recombinant CheD to test binding directly. *B. subtilis* CheD behaved well and could be produced at high levels. However, no interaction between recombinant CheD and SwrD was observed by affinity interaction studies or SEC (Fig. S5). We then overexpressed full-length MotA/B using established protocols (*41*), but again could not detect an interaction of the complex with SwrD, using either affinity tagged MotB or affinity tagged SwrD as “bait” (Fig. S6). In the first case, experiments were limited by low amounts MotA/B and non-specific binding of SwrD to the affinity resin (Fig. S6A), although in the second case, large amounts of SwrD did not recruit overexpressed MotA/B from solubilized lysate (Fig. S6B). Thus, the high confidence AlphaFold predictions of SwrD interaction partners do not seem to bear out in these cases, perhaps suggesting that other factors may be involved in these interactions.

## Conclusion

Herein, we have determined structures and solution properties of the swarm-influencing protein, SwrD, from *B. subtilis* and *B. burgdorferi*. SwrD forms a hexameric state in solution with a charged, disordered C-terminus and a central pore. Sequence and structural homology indicates that SwrD shares similarity to diverse proteins that includes the YgdR family and various gene regulators, but lacks the structural elements these proteins use to either undergo secretion and lipidation or recognize DNA/RNA. Common to these proteins is ring structure anchored by specific conserved residues in the central pore. The *B. subtilis* 3A38 SwrD k/o strain was devoid of swarming behavior and showed significant dysregulation of *sigD* and *flgM*, which may be a response to the observed swarming defect. Computational interaction assays of SwrD identified potential motility (MotA) and chemotaxis proteins (CheD), but neither could be biochemically validated, as of yet. Nonetheless, the hexameric structure of SwrD is likely key to its function because the most conserved residues in the protein either maintain the hexameric subunit interfaces or situate on the hexamer periphery, where they are available for molecular interactions.

## Methods

### Cloning, expression and purification

Bb SwrD was sub-cloned from a pQE-30 backbone, into a pet28(+) backbone in frame to an N-terminal 6x His tag and thrombin cuts site. All point mutations were made using the Kinase-Ligase-DpnI method to the pet28-Bb SwrD plasmid with successful clones verified through sanger sequencing. Bs SwrD was acquired from Twist Biosciences in frame to an N-terminal 6x His tag and thrombin cut site.

All proteins were expressed in BL21(DE3), in an identical manner. In brief, transformed BL21(DE3) cells were grown to an OD of 0.6 at 37°C, protein expression was subsequently induced with 1mM IPTG and grown for a further 3 hours at 37°C. The cultures were then centrifuged down at 8000 x g for 10 minutes and the pellets were stored at-80°C until required.

Pellets were resuspended in 50mM Tris,500mM NaCl and 20mM imidazole prior to sonication at 70% amplitude (0.3s on, 0.7s off) for two minutes. The lysate was then clarified through high-speed centrifugation, 100 000 x g for one hour, with the supernatant then applied to a His-trap column. The protein was then eluted using an imidazole gradient, 20mM to 500mM, with peak fractions collected and concentrated to ∼2mL. Eluants were then subjected to size-exclusion using a Superdex-200 (S200) increase column with a 20mM Tris 150mM NaCl (GFB) mobile phase.

Peak fractions were concentrated and subjected to overnight thrombin cleavage at room temperature and rerun over a S200 column with a GFB mobile phase. Peak fractions were concentrated to 10mg/mL, snap frozen and stored at-80°C until further use.

### Size Exclusion Chromatography-Multi-Angular Light Scattering (SEC-MALS)

All SEC-MALS experiments were conducted with an in-line S200 column attached to a static 18-angle light scattering detector (DAWN HELEOS-II) and a refractive index detector (Optilab T-rEX) (Wyatt Technology, Goleta, CA). 200ug of protein was injected per run with GFB as the mobile phase. All samples were processed using ASTRA VI with monomeric fraction of BSA (Sigma, St. Louis, MO) used to standardize the light-scattering detectors and maintain data quality control.

### Protein crystallization

All variants were crystalized using the sitting drop method. For protein concentrations, crystal conditions and cryogen conditions see table SX. Data was collected either at Corne High Energy Synchrotron Source (CHESS) or the Advanced Photon Source (APS), denoted in table SX. All diffraction images were indexed and integrated into HKL-2000. Modeling and real-space refinement were carried out in Phenix and Coot, respectively, with a truncated (1–50) AlphaFold model (AF-O51281-F1) used as the initial model for Bb SwrD. All other structures were phased against the experimentally determined Bb SwrD structure. For the last pass refinement, models and experimentally determined densities were run through the PDB-REDO server (*42*).

### *B. subtilis* 3A38 growth and strain construction

*B. subtilis 3A38* was acquired from the Bacillus Genetic Stock Center. For general growth, *B. subtilis* 3A38 was grown in Luria Broth (LB) at 37°C. We utilized homologous repair with an erythromycin cassette to replace swrD, swrD::erm, under the native promoter in a pTwist cloning vector. Briefly, *B. subtilis* 3A38 was grown to an OD ∼0.8 in a modified Spizizen’s competency media (MC, in 100mL and composed of: 1.401 g K_2_HPO_4_.3 H_2_0, 0.54 g KH_2_PO_4_, 2 g Glucose, 1 ml trisodium citrate (300 mM), 0.1 ml ferric ammonium citrate (22 mg/ml), 0.1 g casein hydrolysate and 0.2 g potassium glutamate), prepared fresh and filtered using a 0.22μM syringe filter (*43*, *44*). The culture was then incubated with 500ng of plasmid DNA for two hours prior to plating on LB plates with 1ng/μL of erythromycin and 25ng/ μL lincomycin. Successful transformants were analyzed using sanger sequencing to confirm knock out.

### Swarm expansion assay

Swarm expansion assays were adapted from Philps et al. with minor modifications (*45*). Briefly, swarm plates consisted of 20 mL of LB supplemented with 0.7% agar and air dried in a fume for 20 minutes. WT and *ΔswrD* cultures were grown in LB, pelleted and resuspended in Phosphate Buffer Saline (PBS) to an OD of 1.0, with 10μL subsequently spotted on the swarm plate and left to air dried for 20 mins under septic conditions. Plates were then incubated at 37°C and 50% humidity for 8 hours.

### Real-Time Quantitative PCR

WT and Knockout strains were grown LB at 37°C to an OD 1.0. Pellet was collected through centrifugation at 5000 x g for 10 minutes. RNA was extracted using RNeasy Mini Kit (Qiagen). Contaminating gDNA was removed through TURBO DNA-*free*™ Kit (Invitrogen) and cDNA was produced using High-Capacity cDNA Reverse Transcription Kit (Thermo Fisher). qPCR was conducted using 2× Universal SYBR Green Fast qPCR Mix (Abclonal) with QuantStudio™ 7 Flex Real-Time PCR System. Relative gene expression was calculated using the ΔΔCq method with normalization to *gyrA*.

### AlphaPulldown and Alphafold3

AlphaPulldown was utilized with the ColabFold database, with the bait sequence being WT B.subtilis SwrD and the candidates being all B.subtilis swarming proteins as defined by SubtlWiki(*39*, *46*). Successful candidates were then run through the AlphaFold3 server according to the relevant oligomeric states (*47*).

### MotA/MotB and CheD expression, purification and recombinant pull-downs

To test whether the stator (MotA + MotB) and/or CheD interacts with SwrD we employed recombinant pulldowns using a C-terminally Twin-Strep (TWS) tagged Bs MotB, co-expressed and purified with Bs MotA as previously described (*41*). CheD was N-terminally TWS tagged and expressed using BL21(DE3) in the same manner as SwrD, apart from affinity chromatography modified to follow manufacturers procedure for Strep-Tactin resin (IBA Life Science).

To test the potential SwrD – Stator interaction, Strep-Tactin resin was functionalized with 3-fold excess of the Bs stator. The resin was then washed with 5-fold excess stator binding buffer (50mM Tris, 150mM NaCl, 0.01% LMNG). The resin was then loaded with 3-fold excess SwrD and allowed to incubate at 4°C with gentle agitation for 1 hour. The flow-through was discarded and the resin was then washed twice with 10-fold stator binding buffer, relative to resin volume. Resin was then loaded on the SDS-PAGE gel for direct analysis.

We performed the reverse pulldown utilizing MotA/B expressed in the same manner as previously described. MotA/B were solubilized using the detergent based Bug Buster (Millipore). Ni-resin was saturated with 6X-His tagged SwrD and allowed to incubate with solubilized lysate containing MotA/B. A control sample was prepared by using Ni-resin lacking SwrD. Samples were washed in 10 fold excess of resin volume twice and subsequently eluted with 200 mM imidazole.

To test the potential SwrD - CheD interaction, SwrD and CheD were mixed in a 1:1 stoichiometry at 4°C with gentle agitation for 1 hour. The sample was loaded in 3-fold excess onto Strep-tactin resin and allowed to incubate at 4°C with gentle agitation for 1 hour. The flow-through was discarded and the resin was then washed twice with 10-fold excess, relative to resin volume, with binding buffer (50mM Tris, 150mM NaCl). The samples were then eluted with binding buffer supplemented with 100mM Biotin. The resin was then eluted with 1X sample buffer. A control reaction was performed in the same manner, in the absence of CheD.

## Acknowledgements

This work supported by NIH grants R35GM122535 to BRC and AI078958 to CL. We thank Dr Anikita Sachala and Professor John Helmann for thoughtful discussions on *Bacillus subtilis* culturing and aiding in genetic engineering. Computing was performed with assistance from Cornell Bio-High Performance Computing facility (BIOHPC). X-ray crystal diffraction data was made possible due to the operation and expertise of the Cornell High Energy Synchrotron Source (CHESS) and the Advancement Photon Source (APS).

## Data Accessibility

The crystal structures of *B. subtilis* SwrD and *B. burgdorferi* SwrD have been deposited in the PDB under codes: 36DV, 36DR, 36DO, 36DQ and 36DT.

The authors declare no competing interests.

Abbreviations and symbols: Meaning / use in document
Bb: Borrelia burgdorferi
Bs: Bacillus subtilis
CTT: C-terminal tail
FM / FMs: Flagellar motor(s)
HTH: Helix-turn-helix
k/o: Knock out gene deletion
SEC-MALS: Size-exclusion chromatography–multi-angle light scattering
TWS: Twin-Strep tag
ΔswrD: swrD deletion mutant

