## Supplemental for "The bacterial swarming factor SwrD belongs to a diverse family of proteins that form hexameric rings"

**Supplementary Tables**

**Table S1:** Crystallization and cryogen conditions.

**Table S2:** Data collection and refinement statistics.

**Table S3:** qPCR Primers.

**Supplementary Figures**

**Figure S1:** Disorder prediction of all annotated SwrDs.

**Figure S2:** Sequence alignment of Bb SwrD (top) and Bs SwrD (bottom), with 46% sequence identity and 60% sequence similarity.

**Figure S3:** SwrD purification and Circular Dichroism.(A) Bb SwrD (B) Bs SwrD.

**Figure S4:** SEC-MALS traces and molecular weights. (A)Bb SwrD WT (B) Bs SwrD (WT) (C) Bb SwrD Y17W (D) Bb SwrD Y17R.

**Figure S5:** SDS-PAGE gel of recombinant CheD – SwrD interactions studies.

**Figure S6:** SDS-PAGE gels of recombinant BS stator – SwrD interaction studies.

| Variant | Protein Concentration (mg/mL) | Crystallization Condition | Cryogen |
|---|---|---|---|
| Bb SwrD | 10 | 20%w/v Polyethyleneglycol monomethylether 2,000<br>100mM TRIS; pH7.0 | None (Room Temp diffraction) |
| Bs SwrD | 5 | 0.1M Imidazole, 0.1M Lithium Sulfate, 5% Ethanol | 10% Glycerol |
| Bb SwrD Y17R | 10 | 30% (w/v) PEG 8000,100 mM Sodium acetate/ Acetic acid pH 4.5, 200 mM Lithium sulfate | 20% Ethylene Glycol |
| Bb SwrD Y17W | 10 | 20% (w/v) PEG 1000 100 mM Imidazole/ Hydrochloric acid pH 8.0 200 mM Calcium acetate | 20% Ethylene Glycol |
| Bb SwrD PY1617HH | 10 | 30% PEG 8000<br>200 mM ammonium sulfate | 20% Ethylene Glycol |

**Table S1.** Crystal and cryogen, if used, conditions.

| | Bb SwrD | Bs SwrD | Bb SwrD Y17W | Bb SwrD Y17R | Bb SwrD P16H/Y17H |
|---|---|---|---|---|---|
| **Wavelength** | 0.968 | 1.03 | 1.03 | 1.03 | 0.968 |
| **Resolution range** | 30.0 - 1.8 (1.86 - 1.8) | 44.5 - 2.50 (2.56 - 2.5) | 40.6 - 1.80 (1.84 - 1.8) | 40.6 - 2.09 (2.14 - 2.08) | 49.00 - 2.5 (2.69 - 2.5) |
| **Space group** | P 21 21 21 | P 1 | P 21 21 21 | P 21 21 21 | I 41 2 2 |
| **Unit cell** | 68.0 72.1 75.4 90 90 90 | 61.3 61.2 79.8 98.1 94.8 120.0 | 67.0 71.5 72.9 90 90 90 | 67.2 71.3 72.7 90 90 90 | 98.0 98.0 76.5 90 90 90 |
| **Total reflections** | 433461 | 303831 | 1358358 | 755837 | 1206930 |
| **Unique reflections** | 34546 (2964) | 27564 (1334) | 32924 (2241) | 21235 (1421) | 6502 (1217) |
| **Multiplicity** | 12.4 (11.3) | 3.4 (3.2) | 10.7 (10.6) | 12.2 (12.9) | 23.8 (23.8) |
| **Completeness (%)** | 98.66 (94.8) | 80.87 (53.8) | 99.25 (96.5) | 98.91 (94.2) | 96.78 (93.3) |
| **Mean I/sigma(I)** | 31.0 (3.7) | 14.9 (5.6) | 43.4 (5.4) | 23.2 (5.1) | 51.3 (3.1) |
| **Wilson B-factor** | 26.72 | 37.22 | 21.59 | 24.85 | 58.49 |
| **R-merge** | 0.166 (0.460) | 0.079 (0.155) | 0.104 (0.453) | 0.156 (0.387) | 0.082 (0.764) |
| **R-meas** | 0.173 (0.482) | 0.093 (0.183) | 0.110 (0.476) | 0.164 (0.403) | 0.084 (0.780) |
| **R-pim** | 0.049 (0.142) | 0.049 (0.096) | 0.035 (0.142) | 0.049 (0.112) | 0.018 (0.159) |
| **CC1/2** | 0.989 (0.949) | 0.980 (0.975) | 0.991 (0.954) | 0.967 (0.969) | 1.007 (0.944) |
| **CC*** | 0.997 (0.987) | 0.995 (0.994) | 0.998 (0.988) | 0.992 (0.992) | 1.002 (0.985) |
| **Reflections used in refinement** | 34546 (2964) | 27564 (1334) | 32924 (2241) | 21235 (1421) | 6502 (1217) |
| **Reflections used for R-free** | 1588 (151) | 1905 (90) | 1983 (127) | 1994 (141) | 648 (119) |
| **R-work** | 0.159 (0.228) | 0.211 (0.285) | 0.179 (0.206) | 0.183 (0.180) | 0.231 (0.258) |
| **R-free** | 0.184 (0.280) | 0.240 (0.287) | 0.205 (0.219) | 0.210 (0.232) | 0.259 (0.318) |
| **Number of non-hydrogen atoms** | 2841 | 5265 | 2780 | 2767 | 1352 |
| **macromolecules** | 2576 | 5112 | 2580 | 2562 | 1330 |
| **ligands** | 0 | 10 | 0 | 0 | 8 |
| **solvent** | 265 | 143 | 200 | 205 | 14 |
| **Protein residues** | 319 | 648 | 318 | 318 | 165 |
| **RMS(bonds)** | 0.005 | 0.018 | 0.023 | 0.012 | 0.009 |
| **RMS(angles)** | 0.86 | 1.58 | 1.68 | 1.24 | 1.18 |
| **Ramachandran favored (%)** | 98.0 | 96.1 | 100 | 97.1 | 99.4 |

| | | | | | |
|---|---|---|---|---|---|
| **Ramachandran allowed (%)** | 1.63 | 3.85 | 0 | 2.94 | 0 |
| **Ramachandran outliers (%)** | 0.33 | 0 | 0 | 0 | 0.63 |
| **Rotamer outliers (%)** | 0 | 1.04 | 0.34 | 0.68 | 0.67 |
| **Clashscore** | 1.89 | 3.54 | 5.29 | 4.16 | 3.66 |
| **Average B-factor** | 35.8 | 42.9 | 27.8 | 31.9 | 66.4 |
| **macromolecules** | 34.2 | 43.1 | 28.2 | 32.1 | 66.5 |
| **ligands** | - | 7.8 | - | - | 74.12 |
| **solvent** | 51.4 | 35.5 | 22.4 | 29.2 | 52.4 |
| **PDB** | 36DV | 36DR | 36DO | 36DQ | 36DT |

**Table S2.** Data collection and refinement statistics.

| **Name** | **Primer** |
|---|---|
| gyrA forward | ACGGCCAACTAGTAGCAGTG |
| gyra reverse | GAGACGCACACCTTGAGTGA |
| motA reverse | AATGGAGGACAGGCACCAAG |
| motA forward | GCAACAAAGGCAGCACTGAT |
| srfAA forward | GATCGCCTCAATCGTTTGGC |
| srfAA reverse | TGCCCTCTGTATCGGAGTGA |
| swrAA forward | TGAGGAATGCGCCAAACATT |
| swrAA reverse | TCCACAGCGTGACATTCGAT |
| sigD forward | CCCCAGCCGGGACTTAAAAT |
| sigD reverse | CGGGCGATACATTCCGAAGA |
| flgM forward | ATGATAAGCAAGCGGTGCAA |
| flgM reverse | ACCCGTTTTCAATTTGCGCT |

**Table S3:** qPCR Primers.

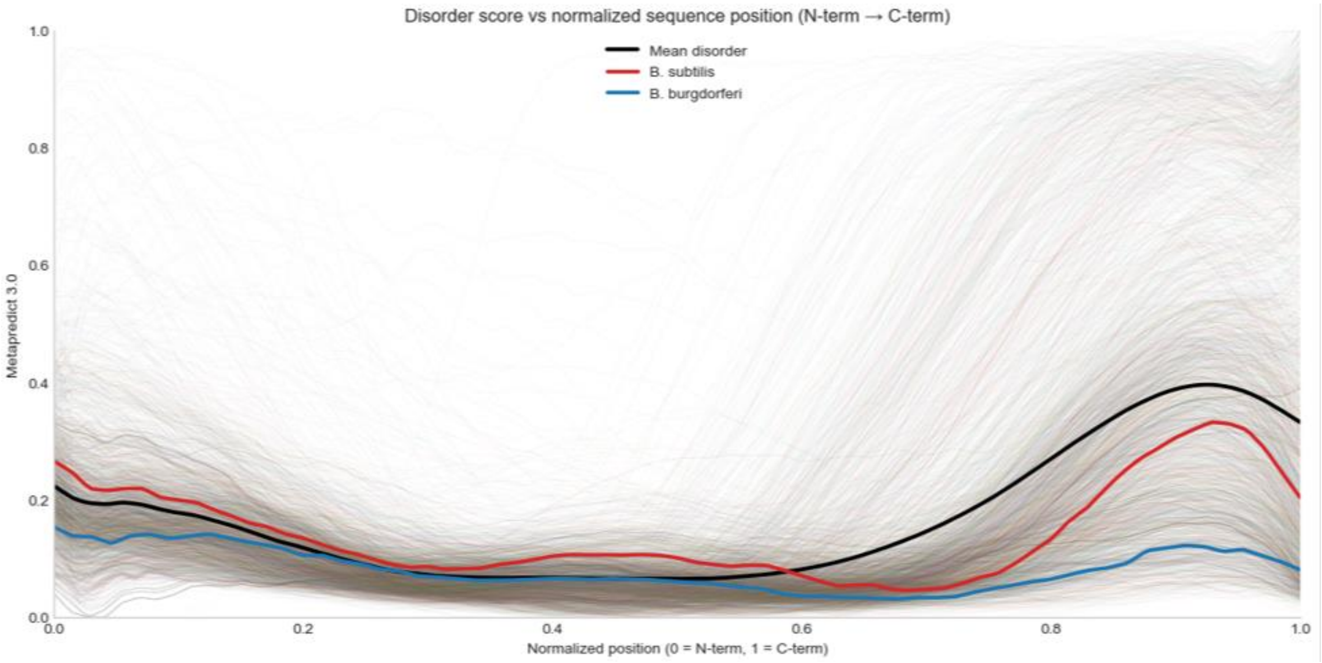


**Fig. S1:** Disorder prediction of all annotated SwrDs.

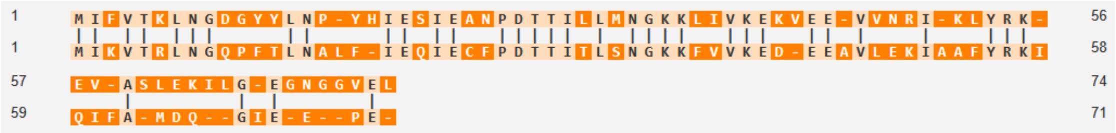


**Fig. S2:** Sequence alignment of Bb SwrD (top) and Bs SwrD (bottom) the proteins have 46% sequence identity and 60% sequence similarity.

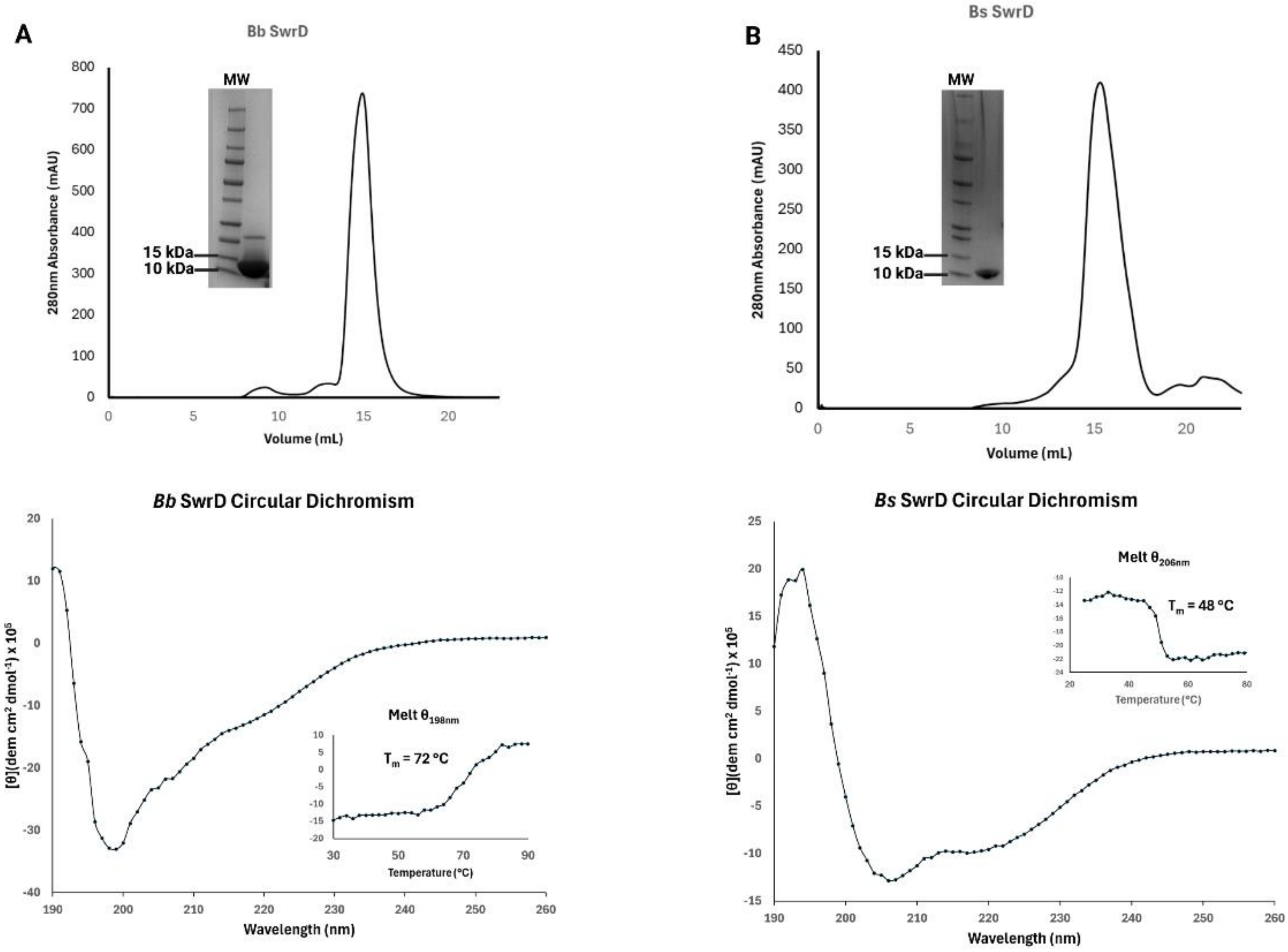


**Fig. S3:** SwrD purification and Circular Dichroism.(A) Bb SwrD (B) Bs SwrD.

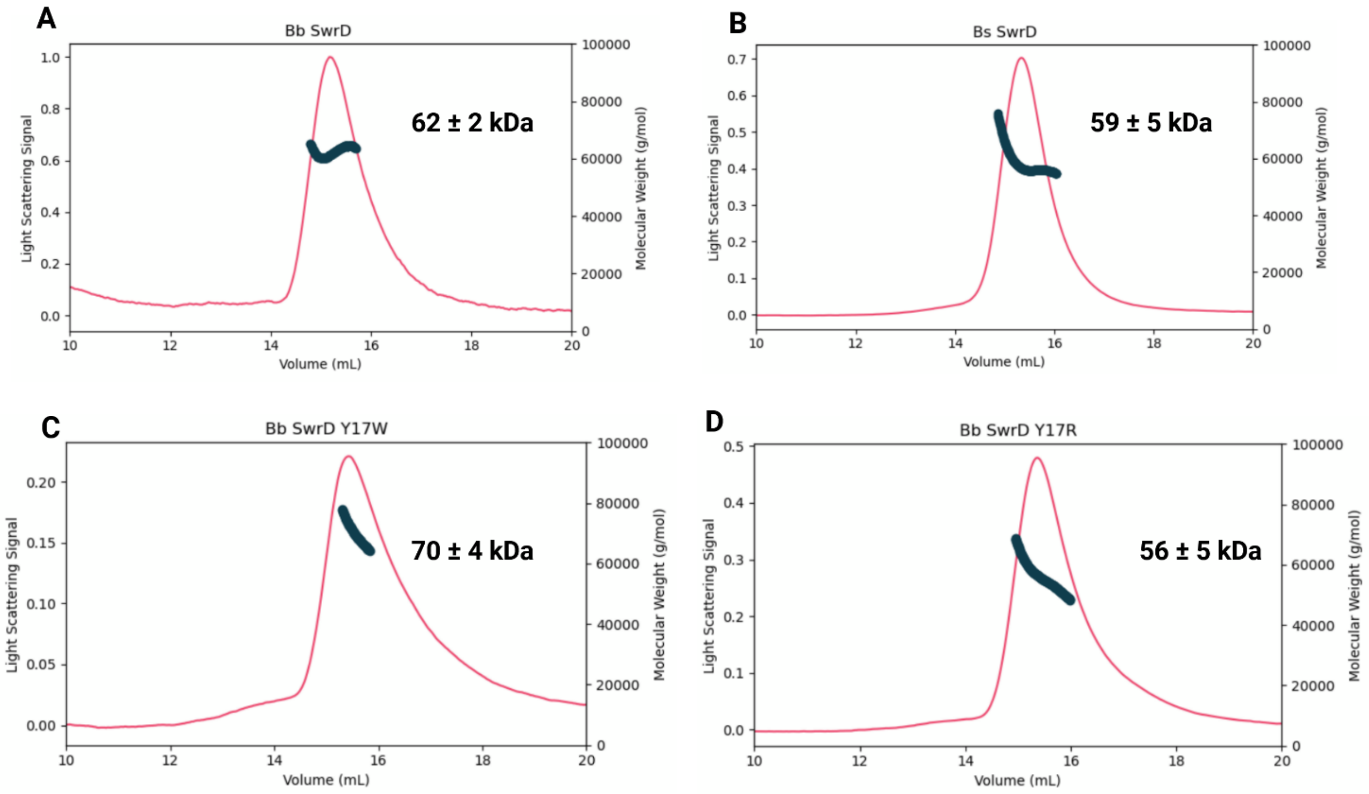


**Fig. S4:** SEC-MALS traces and molecular weights. (A)Bb SwrD WT (B) Bs SwrD (WT) (C) Bb SwrD Y17W (D) Bb SwrD Y17R.

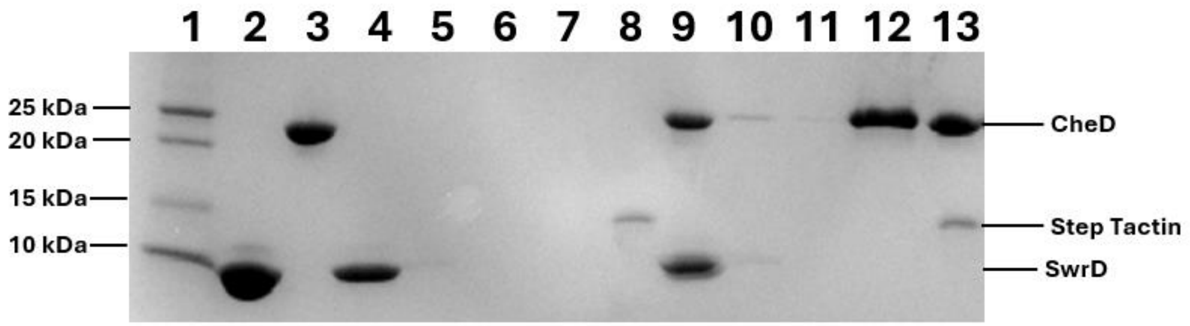


**Fig S5:** SDS-PAGE gel of recombinant interaction studies of strep-tagged CheD with untagged SwrD and control of untagged SwrD binding to the Strep Tactin resin. Lane 1, MW ladder, Lane 2, SwrD Input, Lane 3, CheD input, Lane 4, control flow-through (untagged SwrD alone), Lane 5, control 1st wash 1, Lane 6 control 2nd wash, Lane 7, control elution, Lane 8 control resin (post elution), Lane 9 strep-CheD - SwrD flow-through, Lane 10 strep-CheD - SwrD 1st wash 1, Lane 11 strep-CheD - SwrD 2nd wash, Lane 12 strep-CheD - SwrD elution, Lane 13 strep-CheD - SwrD resin (post elution).

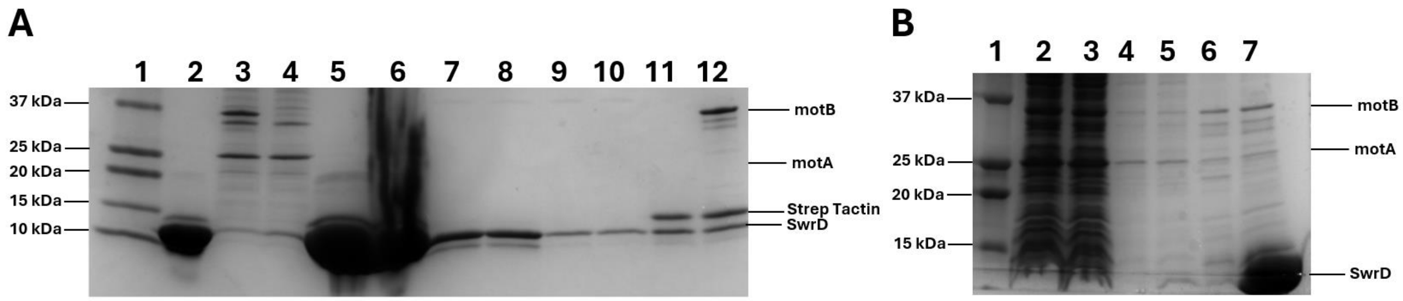


**Fig S6**: Interaction studies of the *B. subtilis* stator (motA/B) and SwrD, performed with either affinity-tagged stator, or affinity-tagged SwrD. (A) MotB is strep-tagged for interaction with Strep Tactin resin, whereas in the "control" lanes unlabeled SwrD is incubated with Strep Tactin resin in the absence of MotA/MotB. Lane 1, Molecular weight ladder (MW), Lane 2, SwrD input, Lane 3, Stator input, Lane 4, Stator loaded resin flow-through, Lane 5, Control resin SwrD flow-through, Lane 6, Stator - SwrD post SwrD incubation flow-through, Lane 7, Control resin 1$^{st}$ wash, Lane 8, stator -SwrD 1$^{st}$ wash, Lane 9, Control 2$^{nd}$ wash, Lane 10, stator-SwrD 2$^{nd}$ wash, Lane 11, control resin and Lane 12, stator- SwrD resin. (B) Detergent solubilized cells expressing motA and motB was incubated with Ni-NTA-resin that was functionalized with (SwrD) or without (Control) His-tagged SwrD. Lane 1, Molecular weight ladder (MW), Lane 2, SwrD loaded resin flow through, Lane 3, Control resin flow-through, Lane 4, SwrD resin Wash, Lane 5, Control resin Wash, Lane 6, Control resin elution with imidazole and Lane 7, SwrD resin elution with imidazole. Neither experiment shows an increase in prey (A, SwrD; B, stator) in the presence of the bait (A, stator; B, SwrD).
